# The diversity of lesion network mapping findings

**DOI:** 10.64898/2026.02.12.705312

**Authors:** Yao Meng, Wenqiang Xu, Jing Jiang, Panpan Hu, Kai Wang, Gong-Jun Ji

## Abstract

Lesion network mapping (LNM) has emerged as a popular framework to map the network mechanism of brain disorders using lesions and normative brain connectome (NBC)^1^. It was first demonstrated in neurological symptoms and was rapidly extended to a broad range of brain disorders in the past decade. A recent study by Van den Heuvel et al. questioned the methodological foundations of LNM^2^. We here raise concerns regarding their errors and biases in the methodology and visualization. The conclusion of that study—LNM maps circumscribed brain changes mostly to one and the same outcome—is not supported by the data presented.

## Results

### Impressive but biased illustrations

Van den Heuvel et al. present an impressive visualization showing apparent similarity between LNM maps and the degree centrality (DC) map across conditions (their Figure 1). However, we noticed that several LNM network patterns clearly diverge from DC yet received minimal presentation. For instance, the PTSD panel (their Figure 1b) displays a pattern dominated by regions that do not align with high-degree hubs. This mismatch to high-degree hubs can also be found in LNM studies of psychosis^3^ and depression^4^, which were not given equal weight in their presentation. Another fascinating illustration involves examples (e.g., agency and autism networks) whose patterns could supposedly be identified using spun lesions or even randomized NBC matrices. Given that psychiatric and neurological disorders often share common network components—as previously demonstrated by Van den Heuvel et al.—some degree of overlap between disease networks is expected^5^. However, this phenomenon does not hold for all LNM networks. A counter-example is the frontotemporal dementia network^6^, whose pattern we cannot be replicated under conditions of spun lesions or randomized NBC matrices (Fig. 1a-c). Besides these biased examples, the full landscape of LNM networks can be glimpsed from the overlapping map (their Figure 1k). According to their theory that LNM networks reflect the DC of the NBC, the top DC areas—specifically in the primary motor and sensory cortices—should be repeatedly identified across LNM studies. However, the overlapping map provides evidence that contradicts this prediction. In contrast, certain lower-DC areas, such as the posterior cingulate cortex, exhibit non-negligible consistency across networks (their Figure 1k).

**Figure 1.**
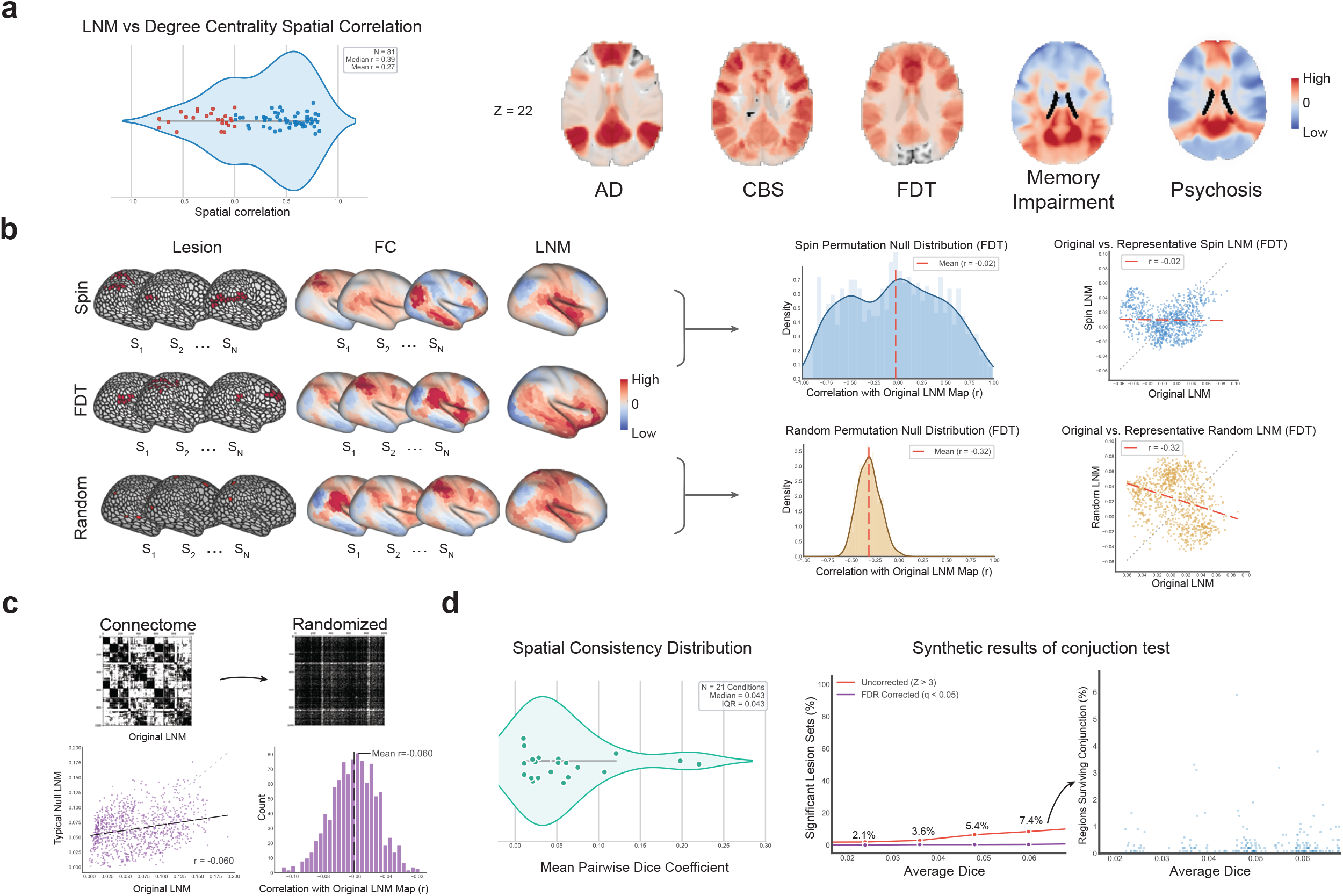
LNM identifies specific network patterns distinct from degree centrality and random null distributions. **a. Dissociation from normative degree centrality.** Left panel showing the distribution of spatial correlation coefficients between condition-specific LNM maps (n=81) and the normative degree centrality map. Red points indicate conditions with non-positive correlations (r ≤ 0), demonstrating that LNM patterns are distinct from generic high-degree hubs. Representative maps (right) illustrate unique topographies for Alzheimer’s Disease (AD), Corticobasal Syndrome (CBS), Frontotemporal Dementia (FDT), Memory Impairment, and Psychosis. **b. Spatial null models demonstrate specific topographic constraints.** Comparison of two null frameworks. Top: The “Spin Null” preserves spatial autocorrelation by rotating lesion centroids (blue). Bottom: The “Random Null” shuffles lesions freely across the cortex (orange). **c. Independence from generic connectome topology.** Verification using degree-preserving randomized connectomes (Maslov-Sneppen rewiring). The scatter plot and histogram display low correlations between original LNM maps and maps derived from randomized networks. **d. Low false-positive rates under realistic lesion heterogeneity.** Left panel: Distribution of mean pairwise Dice coefficients across empirical conditions. Right panel: Simulation results quantifying the rate of lesion sets passed the conjunction test and the percent of significant regions in synthetic cohorts with varying Dice overlap (mean pairwise Dice = 0.024 ~ 0.060, step length = 0.012). FDR correction (purple line) is used in specificity test to further control the false positive rate.

### Errors and biases in methodology

To provide a comprehensive view of the LNM landscape, we reviewed all 102 networks included in the study by van den Heuvel et al. Surprisingly, we identified notable errors and unreliable criteria regarding the collection of LNM networks. 1) Meta-analysis coordinates (https://neurovault.org/collections/GOKZXUGP) were included as LNM findings by mistake^7^; 2) Information from the same study were repeatedly included^8^; 3) The sign direction in three networks were arbitrary inverted; 4) For studies that did not share lesion or prevalence data, the prevalence values—which are often incomplete and inaccurate—were extracted from the figures in the original PDF files; 5) Prevalence maps missed the cerebellum and subcortical parts were also included^9^; 6) Several readily accessible datasets were overlooked, such as the neurodegeneration coordinates from the pioneering coordinate network mapping study^6^ and the rumination coordinates from the initial activation network mapping study^10^. After correcting the LNM network database, we found the mean correlation with DC to be 0.27 (SD = 0.42; 95% CI [0.17, 0.36]; Fig. 1a; Supplementary Table 1). Notably, 23 of the 81 networks exhibited a negative correlation. Furthermore, the original analysis treated multiple networks of the same disorder (e.g., 11 networks for depression) as independent samples, artificially inflating the mean correlation toward a positive value. This corrected landscape reveals high spatial variability across networks, with a substantial portion diverging from the DC distribution. This diversity is consistent with the fundamental principles of LNM deconstructed by van den Heuvel et al. To use an analogy: if the NBC matrix is a dictionary, then pathological changes act as the writer. The writer can compose sentences (networks) in any form or pattern, independent of the frequency with which specific words (nodes) are combined in common usage.

### Ignorance of the difference between networks

Besides the overstatement of similarity between LNM networks, another notable oversight in the work of van den Heuvel et al. is the disregard for inter-network differences. Given that brain areas are highly interconnected and organized into discrete modules or communities, it is no surprise to observe high similarity between certain LNM networks and even healthy control networks. The same phenomenon is common in traditional neuroimaging—whether using seed-to-whole-brain functional connectivity, independent component analysis, or task-based fMRI—where high similarity of whole-brain patterns between healthy and diseased states is inevitable. However, this global similarity is not mutually exclusive with local network divergences, which likely reflect pathology-specific alterations. For instance, Segal et al. identified disease-specific atrophy patterns despite comparable global brain topologies between patients and healthy controls^9^.

### Problems in logic and methodology

Van den Heuvel et al. also addressed the sensitivity and specificity analyses, but several methodological and logical problems warrant further discussion. First, while it is true that one-sample t-tests often produce significant sensitivity results, their utility remains intact. It not only tells us *whether* there is a significant cluster but also show *where* it is. Second, there is a clear logical inconsistency in their argument of specificity analysis. If, as they claim, ‘degree rules all,’ then a specificity analysis comparing real and random networks should logically yield null results. The fact that significant differences emerge in these contrasts strongly supports the view that LNM networks are determined by factors beyond DC. Third, the extent of lesion overlap was likely overestimated in their simulation analysis. While their conjunction analyses showed that the probability of identifying significant LNM maps rose from 10% to 99% as Dice coefficients increased (from 0.08 to 0.3), these parameters lack ecological validity. Empirical data show a mean Dice coefficient of approximately 0.04 (SD = 0.04; Fig. 1d; Supplementary Table 2), suggesting that their simulation relied on unrealistic levels of overlap. We simulated 1000 sets of lesion data with each Dice value (0.024 ~ 0.06, step = 0.012), and found only 7.4% synthetic lesion sets can pass the conjunction test with the highest overlap in the simulation (Fig. 1d).

### Biological significance of LNM

Van den Heuvel et al. further challenged the disease-specific biological significance of LNM using regression analyses, demonstrating that the variance in LNM networks is largely explained by connectome-based matrix features. Crucially, they found that gradient and modular architectures—rather than DC—accounted for the greatest variance. However, this is unsurprising since it is well known that the brain’s functional architecture shapes disease progression which may be captured by LNM networks^11^. The critical gap in our knowledge is identifying which specific matrix properties correspond to the pathophysiological changes of a particular disease. To bridge this gap, recent research has placed a growing emphasis on the functional annotation of these networks^12^. A convincing approach to demonstrate the biological relevance in a given diseases or symptom could be relating the network to clinical treatment. Compellingly, the LNM networks in depression can specifically account for the outcome variance of depression symptom in clinical treatment^13^ and enhanced therapeutic efficacy by identifying symptom-specific targets for precision neuromodulation^14^.

## Conclusion

A primary operational advantage of the LNM framework is its reliance on NBC rather than individual functional imaging. While the work of van den Heuvel et al. serves as a timely opportunity to review the LNM method, their conclusion was biased by methodological flaws. The use of NBC does not prevent us from identifying the disease-specific network mechanisms.

## Methods

**See the methods in Supplementary Materials.**

## Supporting information

Suppementary Methods and Table

## Acknowledgements

This study was funded by the National Natural Science Foundation of China (82371507 to G.J.; U23A20424 and 82090034 to K.W.; 82171917 and 82471271 to P.H.); Distinguished Youth Foundation of Anhui Province (2024AH020004 to G.J.); Postgraduate Academic Innovation Program of Anhui Province (2024xscx058 to W.X.); the Postgraduate Innovation Research and Practice Program of Anhui Medical University (YJS20230062); the Anhui Province Clinical Medical Research Transformation Special Project (202204295107020006 and 202204295107020028 to K.W.); and the Improvement Plan of Scientific Research Platform Foundation Construction of Anhui Medical University (2020xkjT057 to K.W.).

## Author contributions

This text arose out of joint discussions and contributions from all authors.

## Competing interests

The authors declare no competing interests.

## Additional information

Correspondence and requests for materials should be addressed to Gong-Jun Ji.

## Notes

### Competing Interest Statement

The authors have declared no competing interest.

