## Supplementary material for "The diversity of lesion network mapping findings": Suppementary Methods and Table

### Supplementary Methods

#### Spin and random permutation null model

To rigorously validate the specificity of Lesion Network Mapping (LNM) results, we employed two distinct permutation frameworks, illustrated using Frontotemporal Dementia (FDT) as a representative case. Spin Null (Lesion Spin): This method accounts for the inherent spatial autocorrelation of lesion locations. We projected the centroids of empirical lesions onto a spherical surface and applied 1,000 random rotations. The rotational constraint preserves the relative spatial configuration among lesions (i.e., if two lesions are spatially clustered, they remain clustered in the null). These rotated centroids were mapped back to the nearest cortical parcels to generate "Spin Lesion" sets. LNM maps were recomputed for each permuted set to build a null distribution of spatial correlation coefficients. This approach provides a conservative test of whether the results are driven by specific lesion locations rather than general spatial biases. Random Null: As a baseline non-spatial control, we generated null sets by shuffling lesion locations randomly across all cortical parcels, ignoring spatial structure. Comparing the empirical results against this distribution verifies that the observed patterns are not obtainable by chance placement of lesions.

#### Randomized connectome null simulation

To determine if our findings were driven by generic topological properties of the brain graph (e.g., node degree) rather than specific connectivity patterns, we constructed null connectomes using a degree-preserving randomization algorithm (Maslov-Sneppen rewiring). We generated 1,000 randomized adjacency matrices where edges were rewired while maintaining the exact degree of every node. LNM maps were computed using the empirical lesions but with these randomized networks. The low correlation between empirical LNM maps and those derived from randomized connectomes confirms that our results depend on the specific wiring architecture of the human brain.

#### Synthetic lesion conjunction test

We performed a synthetic validation analysis to optimize statistical thresholds for identifying common network substrates. We simulated 1,000 synthetic patient lesion sets (each N=50) with varying degrees of inter-subject lesion overlap, parameterized by the mean pairwise Dice coefficient (range: 0.024–0.06, step length = 0.012).

For each cohort, we applied a dual-threshold conjunction analysis:

Sensitivity Test: We identified regions with strong functional connectivity to the lesion sites by applying a one-sample t-test threshold ($|T| > 7$, corresponding to a large effect size) to each lesion's connectivity map. A region was retained only if it exceeded this threshold in at least 75% of the lesions in the cohort.

Specificity Test: To ensure findings were specific to lesion locations, we generated 1,000 spin-permuted null cohorts for each synthetic dataset. For each brain region, we computed a z-score comparing the observed group-mean connectivity against the null distribution. We applied a standard threshold of $|Z| > 3$ ($p < 0.001$ uncorrected) and compared it against a False Discovery Rate (FDR) corrected threshold ($q < 0.05$).

A region was considered a significant "conjunction" finding only if it passed both the Sensitivity criteria and the Specificity criteria. We calculated the percentage of cohorts producing any significant findings under null-like condition, and the detection yield across the simulated range of heterogeneity. Results demonstrated that FDR correction effectively controls the FPR while maintaining sensitivity across the empirical range of lesion overlap.

### Supplementary Tables

**Supplementary Table 1 Correlations of LNM maps with degree of NBC**

| Map name | Condition | Correlation (r) with degree |
|---|---|---|
| DARBY_DELUSION | Delusional misidentifications | 0.65 |
| ABTEMAN_CONFABULATION | Confabulation | -0.12 |
| BOES_HALLUCINOSIS | Peduncular hallucinosis | 0.13 |
| KUTSCHE_APHANTASIA | Aphantasia | 0.39 |
| BURKE_MIGRAINE | Migraine | 0.7 |
| COHEN_PROSOPAGNOSIA | Prosopagnosia | 0.34 |
| HORN_DYSTONIAGENERALIZEDDBS | Dystonia (generalized) | -0.1 |
| CORP_DYSTONIA | Dystonia (cervical) | 0 |
| ALFATLY_OCULOGYRIC | Oculogyric crises | 0.3 |
| DARBY_AGENCY | Disrupted agency | 0.59 |
| DARBY_CRIMINALITY | Criminality | -0.4 |
| DARBY_VOLITION | Disrupted volition | 0.05 |
| FISCHER_COMACAUSE | Coma (cause) | -0.08 |
| FRIEDRICH_AIWS | Alice in Wonderland Syndrome | 0.36 |
| HAQUE_ALCOHOL | Alcohol use disorder remission | -0.36 |
| HERBET_AWARENESS | Bodily awareness | -0.47 |
| HIGASHIYAMA_ACCENT | Foreign accent syndrome | 0.78 |
| JOUTSA_ADDICTIONCONTRASTAB | Smoking addiction (contrast: A-B) | 0.63 |
| JOUTSA_ADDICTIONGROUPA | Smoking addiction (A, remission for smoking) | 0.81 |
| JOUTSA_ADDICTIONGROUPB | Smoking addiction (B, not quit smoking) | 0.7 |
| JOUTSA_ADDICTIONGROUPC | Smoking addiction (C, quit but did not remit) | 0.82 |
| KLETENIK_ANOSOGNOSIA | Anosognosia (sLNM) | -0.36 |
| KLETENIK_BLINDSIGHT | Blindsight | 0.28 |
| KUTSCHE_CREATIVITY | Creativity (coordinates) | 0.58 |
| LAGANIERE_HEMICHOREA | Hemichorea-hemiballismus | 0.61 |

| | | |
|---|---|---|
| LEE_MANIA | Mania | 0.59 |
| LESIONBANK_HYPERSOMNIA | Hypersomnia | 0.41 |
| LESIONBANK_INSOMNIA | Insomnia | 0.49 |
| LESIONBANK_NEGLECT | Neglect syndrome | 0.7 |
| LI_VERTIGO | Vertigo | 0.78 |
| LIESMAKI_ATAXIA | Ataxia | 0.51 |
| SUN_EPILEPSY | Epilepsy | -0.23 |
| JI_EPILEPSY | Epilepsy (LNM) | 0.66 |
| MANSOURI_EPILEPSY | Epilepsy | 0.39 |
| MOLLICA_PSAC+ | Persisting symptoms after concussion (high) | 0.5 |
| MOLLICA_PSAC- | Persisting symptoms after concussion (low) | 0 |
| PINES_PSYCHOSIS | Psychosis (LNM) | -0.52 |
| MARCELINO_PALSY | Dyskinetic cerebral palsy | 0.6 |
| QIN_SPASTICITY | Spasticity | 0.68 |
| REICH_PARKINSONDBS | Parkinson's disease (cognitive decline) | -0.64 |
| FASANO_GAIT | Freezing gait | 0.44 |
| JOUTSA_HOLMES | Holmes tremor | 0.01 |
| JOUTSA_TREMOR | Tremor (sLNM) | 0.7 |
| JOUTSA_PARKINSONISM | Parkinsonism | 0.29 |
| RIOS_ALZHEIMERDBS | Alzheimer's disease | -0.08 |
| SIDDIQI_POLITICAL | Political involvement | 0.46 |
| SIDDIQI_PTSD | PTSD (sLNM) | -0.27 |
| STUBB_SUD | Substance abuse disorder | 0.21 |
| TAYLOR_TRANSDIAGNOSTIC | Transdiagnostic psychiatric disorders | 0.14 |
| THEYS_STUTTERING | Neurogenic stuttering | 0.66 |
| WANG_AMNESIA | Transient global amnesia | -0.21 |
| FERGUSON_AMNESIA | Amnesia | -0.12 |
| KLETENIK_MS | MS (memory impairment) | -0.04 |
| YUAN_APNEA | Apnea | -0.09 |

| | | |
|---|---|---|
| YUAN_RBD | Rapid eye movement sleep behavior disorder | -0.06 |
| ZOUKI_TICS | Tics (Tourette syndrome) | 0.34 |
| GANOS_TICS | Tics | 0.53 |
| GERMANN_OCDCAUSE | OCD (cause) | 0.42 |
| LI_OCDDBS | OCD | 0.39 |
| SIDDIQI_DEPRESSION | Depression | 0.28 |
| SIDDIQI_MSDEPRESSION | MS (depression) | 0.44 |
| LI_DEPRESSION | Depression | 0.29 |
| FOX_DLPFCTMS | Depression | 0.53 |
| CASH_DEPRESSIONEMO | Depression (emotional) | 0.74 |
| CASH_DEPRESSIONCOG | Depression (cognitive) | 0.58 |
| JI_ANTIDEPRESSANT | Depression remission | 0.54 |
| TRAPP_DEPRESSIONNEG | Depression (resilience) | -0.13 |
| TRAPP_DEPRESSIONPOS | Depression (risk) | -0.03 |
| SIDDIQI_ANXIETY | Anxiety-depression symptoms | -0.56 |
| GONG_CONTROLFORRUMIN ATION | Facial processing (control rumination) | 0.76 |
| GONG_EMOTION | Facial processing (emotion) | 0.77 |
| GONG_NONEMOTION | Facial processing (non-emotion) | 0.75 |
| ZARIFKAR_APHASIA | Aphasia | 0.74 |
| SIHVONEN_AMUSIA | Amusia | 0.8 |
| SOUTER_APHASIA | Aphasia (semantic) | 0.44 |
| ZARIFKAR_APRAXIA | Apraxia (eye-opening) | 0.78 |
| Jiang_EmoRegulation[1] | Emotion regulation | 0.01 |
| Peng_ANM[2] | Rumination | -0.73 |
| Darby_AD[3] | AD | -0.73 |
| Darby_CSB[3] | CSB | 0.08 |
| Darby_FDT[3] | FDT | -0.16 |

**Supplementary Table 2 Mean pairwise Dice across conditions with avialable lesion data from ven de Heuvel et al.**

| Condition | N_Lesions | Mean Pairwise Dice |
|---|---|---|
| ASD | 9 | 0.01 |
| Anxiety | 16 | 0.02 |
| BD | 29 | 0.02 |
| MDD | 16 | 0.06 |
| OCD | 13 | 0.01 |
| PTSD | 16 | 0.06 |
| SCZ | 159 | 0.03 |
| ADDICTION | 30 | 0.05 |
| AGENCY | 45 | 0.02 |
| APHASIA_RECOVERY | 217 | 0.22 |
| AliceInWonderland | 25 | 0.03 |
| Amnesia | 44 | 0.08 |
| EPILEPSY | 111 | 0.01 |
| LESYMAP | 131 | 0.20 |
| MDDDLPFC_TMS | 20 | 0.12 |
| MIGRAINE | 8 | 0.11 |
| NEGLECT | 27 | 0.04 |
| OCD | 100 | 0.01 |
| PARKINSONISM | 18 | 0.06 |
| SCHIZOPHRENIA | 100 | 0.01 |
| STUTTERING | 14 | 0.05 |
